# Transcriptome analysis of *Xanthomonas oryzae* pv. *oryzicola* exposed to H_2_O_2_ reveals horizontal gene transfer contributes to its oxidative stress response

**DOI:** 10.1101/669614

**Authors:** Yuan Fang, Haoye Wang, Xia Liu, Yuchun Rao, Xin Dedong, Bo Zhu

## Abstract

*Xanthomonas oryzae* pv. *oryzicola* (*Xoc*) is the causal agent of bacterial leaf streak in rice. It is known as one of the most severe seed-born bacterial diseases of rice, molecular role governing its interaction with rice is mostly still unexplored. To successfully invade rice, the survival of the *Xoc* is mandotarory following generating a specific response to its host’s oxidative stress condition. However, the response network of *Xoc* is still unknown. To address this question, we performed a time-series RNA-seq analysis on the *Xoc* response to H_2_O_2_. Overall, our RNA sequence analysis of *Xoc* revealed a significant global gene expression profile when it exposed to H_2_O_2_. The response of key genes was also noted that soxR triggers and regulates the *Xoc* oxidative stress response in the early stage of infection, gene expression kinetics among the time-series samples, namely for TonB-dependent receptors and the *suf* and *pst* operons. Moreover, a hypothetical protein (XOC_2868) showed significant differential expression following its mutant endorsed RNA-seq findings by clearly displaying a greater H_2_O_2_ sensitivity and decreased pathogenicity than the wild-type. Gene location and phylogeny analysis strongly suggests that this gene may have been horizontally transferred from a *Burkholderiaceae* ancestor. Our study not only provides a first glance of *Xoc*’s global response against oxidative stress, but it also reveals the impact of horizontal gene transfer in the evolutionary history of *Xoc*.

## Introduction

Oxidative burst is a process in which high concentrations of reactive oxygen species (ROS) are produced at the plasma membrane in the vicinity of a pathogen [1, 2]. This plant activity directly kills the pathogen or slows its growth by producing toxins or by acting as a signaling cascade which may lead to multifarious defenses, including the hypersensitive response (HR) and cell wall modifications [3]. Since ROS activity is a common feature of plant defense systems and the mechanism of pathogen cell death, any pathogen able to resist its effect is likely advantaged. Although this process, of overcoming ROS, is critical for a phytopathogen to successfully invade a host plant, until now no systematic time-series response research has yet been carried out on any phytobacteria.

RNA-seq, a relatively new transcriptomic analysis method, has been widely used to investigate both gene expression and content in many bacteria [4]. Much of the oxidative stress research done on human pathogens and environmental bacteria has been carried out with this technology: for example, the studies *Salmonella enterica* and *Ralstonia eutropha* [5, 6]. However, the application of RNA-seq to phytobacteria remains very limited, with no such research yet done on *Xanthomonas oryzae* pv. *oryzicola* (*Xoc*) [7].

*Xanthom*onas is among the most widely distributed genus of Gram-negative bacteria that have the typical characteristic of yellow pigment. This genus consists of a number of phytopathogenic bacteria that infect more than 400 host species, including a wide variety of economically important plants, such as rice, citrus, banana, cabbage, and bean [8, 9]. One species of this genus, *Xoc*, is the causal agent of bacterial leaf streak (BLS) in rice (*Oryza sativa*). BLS is considered one of the most potent seed-born bacterial diseases of rice in many Asian countries and parts of Africa [10]. The interaction between *Xoc* and rice is complex, however. This complexity is the outcome of a longstanding and ongoing evolutionary battle, in which the bacteria attempts to invade and multiply, while the rice cell attempts to recognize and defend itself from this invasion. Of these plant responses, oxidative burst is thought to be one of the frontier response against pathogens [2].

BLS256, the representative genome of *Xoc*, has been whole-genome sequenced [11]. Interestingly, more than 30% of the coding genes in this genome are hypothetical gene. Indeed, to date, very few molecular investigations of *Xoc* have been carried out. In this context, proteomics data can greatly help to clarify and solve the puzzle behind these genes. For example, by using protomics data, Ram et al. uncovered many new genes, previous annotated as hypothetical proteins, which had important roles to play in biofilm formation [12].

In this paper, the Illumina RNA-Seq platform was used to identify genes differentially expressed by the *Xoc* strain BLS256 in response to a time-series H_2_O_2_ treatment (early, middle, and late). Besides the genes previously confirmed to be triggered by oxidative stress, we found one hypothetical protein: it showed a high-fold differential expression compared with the control following deep bioinformatics analysis strongly suggested that this gene could have been horizontally transferred from other organisms.

## Materials and Methods

### Strains, plasmids and culture conditions

The bacterial strains and plasmids used in this study are listed in S1 Table. *Escherichia coli* strains were cultivated at 37 °C in Luria-Bertani (LB) medium or on LB agar plates. Unless specified otherwise, *Xoc* strains were grown at 28 °C in NB (0.1% yeast extract, 0.3% beef extract, 0.5% polypeptone, and 1% sucrose), NA (NB with 15 g L^−1^ agar), NAN (NA without sucrose), NAS (NA with 10% sucrose), and NY (NB without beef extract and sucrose). In some experiments, strains were grown in MMX minimal medium [0.5% glucose, 0.2% (NH_4_)_2_SO_4_, 0.1% trisodium, citrate dihydrate, 0.4% K_2_HPO_4_, 0.6% KH_2_PO_4_, 0.02% MgSO_4_ • 7H_2_O]. Antibiotics were added at the following concentrations (μg mL^−1^) when required: kanamycin (Km), 50; ampicillin (Amp), 100.

### Oxidative stress treatments and total RNA preparation

H_2_O_2_ resistance assays were performed as described previously [13] with some modifications. *Xoc* strains were cultured to the mid-log phase (OD 600 = 1.0 ~ 1.2) in NB medium. Cultures were treated with 0.1mM H_2_O_2_ in a 28 °C shaking incubator. At time of 0, 7, 15, and 45 min, aliquots were withdrawn and pelleted by centrifugation at 4 °C. Cell pellet was washed two times with cold PBS and the total RNA was extracted immediately following the manual of RNeasy Protect Bacteria Mini Kit (QIAGEN). Two biological replicates were performed in this experiment.

### mRNA purification and cDNA synthesis

Ten micrograms from each total RNA sample was treated with the MICROBExpress Bacterial mRNA Enrichment kit and RiboMinus™ Transcriptome Isolation Kit (Bacteria) (Invitrogen). Bacterial mRNAs were chemically fragmented to the size range of 200-250 bp using 1 × fragmentation solution for 2.5 min at 94°C. cDNA was generated according to instructions given in SuperScript Double-Stranded cDNA Synthesis Kit (Invitrogen). Briefly, each mRNA sample was mixed with 100 pmol of random hexamers, incubated at 65°C for 5 min, chilled on ice, mixed with 4 μL of First-Strand Reaction Buffer (Invitrogen), 2 μL of 0.1 M DTT, 1 μL of 10 mM RNase-freed NTPmix, 1 μL of SuperScript III reverse transcriptase (Invitrogen), and incubated at 50°C for 1 h. To generate the second strand, the following Invitrogen reagents were added: 51.5 μL of RNase-free water, 20 μL of second-strand reaction buffer, 2.5 μL of 10 mM RNase-free dNTP mix, 50 U *E. coli* DNA Polymerase, 5 U *E. coli* RNase H, and incubated at 16 °C for 2.5 h.

### Library construction and Illumina sequencing

The Illumina Paired End Sample Prep kit was used for RNA-Seq library creation according to the manufacturer’s instructions as follows: Fragmented cDNA was end-repaired, ligated to Illumina adaptors, and amplified by 18 cycles of PCR. Paired-end 100-bp reads were generated by high-throughput sequencing with the Illumina Hiseq2000 Genome Analyzer instrument.

### RNA-Seq data analysis

After removing the low quality reads and adaptors, pair-end reads were mapped to the reference genome BLS256 by using bowtie1.1.1 with default parameters [14]. If reads mapped to more than one location, only the one showed the highest score was kept. Reads mapping to rRNA and tRNA regions were removed from further analysis. Since four time-points samples were prepared in this study (0min, 7min, 15min and 45min; defined as WT, early, middle and late stage), All the other samples were always compared with the 0 min samples to detect the differential expressed genes (DEGs). After getting the reads number from every sample, edgeR with TMM normalization method was used to determine the DEGs [15]. FDR value < 0.05 was selected as the cutoff for further analysis. Cluster 3.0 and treeview were used to represent the cluster of DEGs over time-series samples [16].

### Quantitative real-time PCR

Bacterial total RNA was extracted as described by the manual of RNeasy Protect Bacteria Mini Kit (QIAGEN) and then was used for generating the first strand complementary DNA (cDNA) as described in the protocol of the Takara PrimeScript RT reagent Kit with gDNA Eraser (Takara). Briefly, 1 mL of bacterial cells were mixed with 2 mL of RNAprotect Bacteria Reagent before incubating for 5 min at room temperature. The pellet was obtained after centrifugation and was then treated by TE buffer (10 mM Tris·Cl, 1 mM EDTA; pH 8.0) containing 1 mg mL^−1^ lysozyme at room temperature for 5 mins. After detecting the RNA quantity and quality by Nanodrop ND1000 spectrophotometer V 3.5.2 (NanoDrop Technologies, Wilmington, DE, USA), 1 μg of the resulted total RNA was incubated at 42 °C for 2 mins to eliminate the gDNA by gDNA Eraser before obtaining cDNA. Then the reverse transcription reaction was accomplished by incubating at 37 °C for 15 min and then 85 °C for 5 s in the presence of random RT primers. The cDNA was then used directly as the template for qRT-PCR using a SYBER Green master mix (Protech Technology Enterprise Co., Ltd.) on an ABI Prism 7000 sequence detection system (Applied Biosystems). Normalized expression levels of the target gene transcripts were calculated relative to the rRNA using the ΔΔCT method, where CT is the threshold cycle. Three biological replicates were carried out in this experiment.

### Construction of defective deletion mutant and complementation

To investigate the role of interested genes in *Xoc*, In-frame deletion mutations were constructed in BLS256 using homologous recombination. Briefly, two fragments flanking the left and right of corresponding genes were amplified from the wide-type genomic DNA with primer pairs listed in S2 Table. The amplified fragments were cloned into **pMD18-T** (TaKaRa), confirmed by sequence analysis, and then digested and subcloned into vector **pKMS1** [17] at *BamH*I and *Pst*I (or *Sal*I) sites. The resulted recombinant plasmids were introduced into BLS256 by electroporation, and transformants were plated on NAN plates supplemented with kanamycin. Colonies resulting from a single homologous crossover (integration of deletion construct at either the left or right border of target gene) were then transferred to NBN broth, grown for 12 h at 28°C, and then plated on NAS plates for sucrose-positive deletion mutant selection. Sucrose resistant colonies were visible within 3 to 4 days and then transferred to NA plates and NA plus kanamycin plates. Since kanamycin-sensitive colonies could be mutants containing a second homologous crossover, these were further examined by PCR amplification with the primer pairs.

In order to complement the deletion mutants, the full-length of corresponding genes including promoter regions were amplified using primer pairs listed in S2 Table. The amplified DNA fragments were cloned into **pUFR034** [17] at the *BamH*I and *Pst*I (or *Sal*I) sites to create the recombinant plasmids. The recombinant plasmids were transferred into corresponding mutants by electroporation, and transformants were screened on NA plates with kanamycin.

### H_2_O_2_ resistance assay

NB agar plates were prepared containing H_2_O_2_ concentrations of 0, 0.1 and 0.25 mM, respectively. Strains were cultured to the mid-log phase (OD 600 = 1.0) in NB medium. 5-μL aliquot of the initial culture and diluted cultures for each strain were spotted onto NB agar plates (in triplicate) and cultured for 36 h at 28 °C [18].

### Pathogenicity assays

Bacteria, which were prepared based on previous reported method [18], were inoculated into leaves of adult rice plants (*Oryza sativa* cv. IR24, susceptible to BB, 2months old) for evaluating water-soaked symptoms. All plants were maintained in a greenhouse as described previously [11]. Plant phenotypes were scored 24 h post-inoculation for the HR in tobacco, 3 days post-inoculation (dpi) for water-soaked symptoms, and 14 dpi for lesion length. Five leaves were inoculated for each independent experiment, and each treatment was repeated at least three times.

### Phylogenetic analysis

Interested gene was first compared against nr database for homologs searching with E<0.0001 as cutoff. Next, 40% alignment similarity and 80% coverage parameter was used to define the real homologue genes. Mafft was used to generate the multiple sequences alignment [19]. Maximum Likelihood phylogeny was finally evaluated and built by PhyML [20] using a JTT model and a gamma distribution with eight rate categories. We performed 1,000 bootstraps to gain branch support values.

## Results

### Global overview of the RNA-Seq data

In this study, 100-bp paired-end deep sequencing was performed on all the tested samples. In general, more than 40M reads were generated from each single sample. After the adaptor removal, quality control, and removal of the reads, which were mapped to the ribosomal RNA, 7M to 10M confident reads remained. The sequencing depth in this experiment was more than 150×, sufficient for further statistical analysis. In general, 7, 177, and 246 genes were differentially regulated in the early, middle, and late stages, respectively (Fig 1, S3 Table). The overall number of differentially expressed genes (DEGs) we found was rather similar to that in recent research of human pathogens and environmental bacteria [21, 22], the exception was DEGs in the early state (7 min), a number almost 10× smaller in our study, which indicates the oxidative response difference that may occur between plant pathogen versus other bacteria. To verify the accuracy of this RNA-seq data, qRT-PCR was carried out on 30 randomly selected candidates. The result from a Pearson correlation coefficient test confirmed that our RNA-seq data was robust and could be trusted (*r* > 0.95, data not shown).

**Fig 1.**
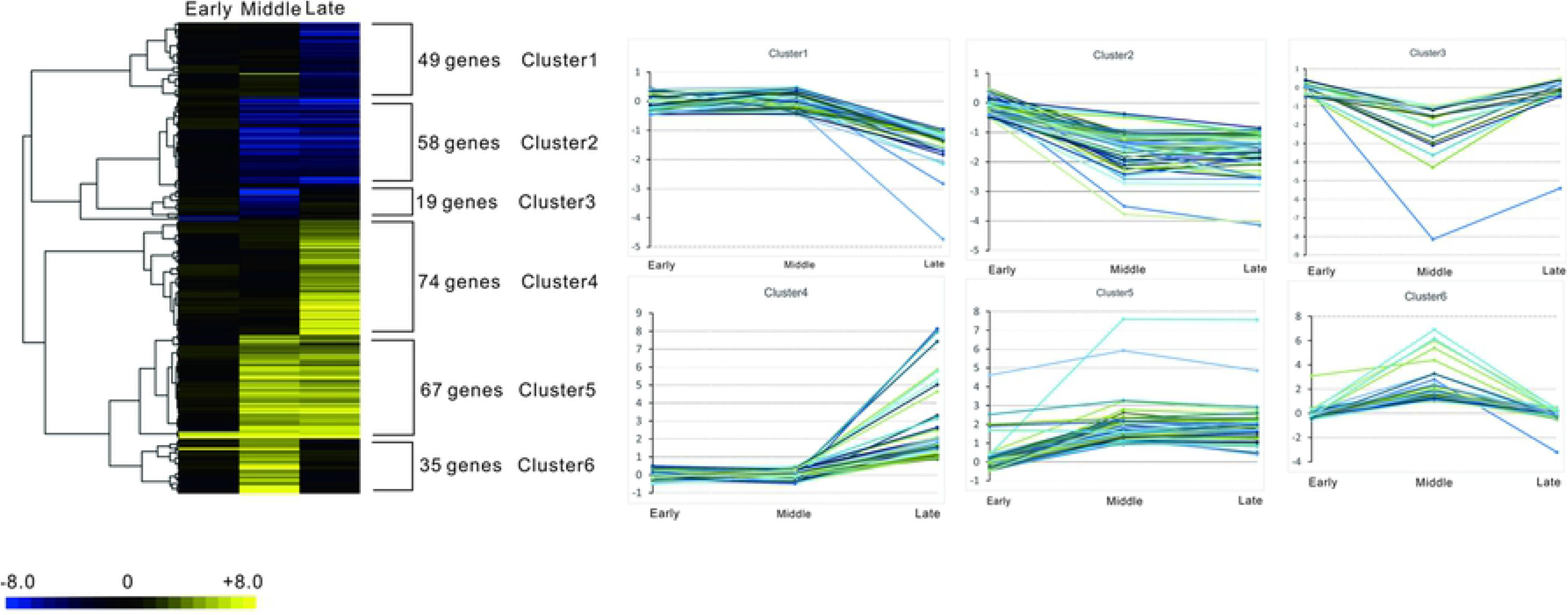
Time-series transcriptome study of *Xoc* (*Xanthomonas oryzae* pv. *oryzicola*) resistance to an oxidative stress response. Hundreds of genes are differentially expressed at different time points (early, middle, and late) during the H_2_O_2_ treatment when compared with the 0-min sample. These differential expressed genes can be classified into different clusters based on their expression patterns. The line from −8.0 to 8.0 represents the Log_2_ fold change value.

### General gene expression kinetics among the time-series samples

Previous research indicates that bacteria require dynamic regulatory networks at different time-points when they are exposed to environmental stress [21, 23]. To capture this variable activity, gene expression among the time-series samples was generated and results are shown in Fig 1. In general, clusters 1 and 2 had a decreased induction whereas clusters 4 and 5 had an increased induction through time (Fig 1). By contrast, the clusters 3 and 6 consisted of those genes whose expression peaked at the middle time stage.

GO and KEGG enrichment analysis was done based on the STRING V10 database [24], with a *p* < 0.01 after a Bonferroni correction [25] set as the cutoff. The GO enrichment analysis revealed transport (GO:0006810), cell outer membrane (GO:0009279), and acyl-CoA dehydrogenase activity (GO:0003995) as significant based on this cutoff for clusters 1 and 2 (Table 1). The regulation of cellular process (GO:0050794), chemotaxis (GO:0006935), single organism signaling (GO:0044700), response to external stimulus (GO:0009605), molecular transducer activity (GO:0060089), and signal transducer activity (GO:0004871) were significant for clusters 4 and 5 (Table 1). Membrane protien (GO:0016020), outer membrane protein (GO:0019867), and cellular component (GO:0005575) were significant for clusters 3 and 6 (Table 1). The KEGG enrichment results suggested that only valine, leucine, and isoleucine degradation was significant for clusters 1 and 2 (Table 1). Interestingly, we found many TonB-dependent receptors (TBDRs) that were differentially expressed in all the three time stages, indicating their importance in oxidative stress (S3 Table).

**Table 1.**
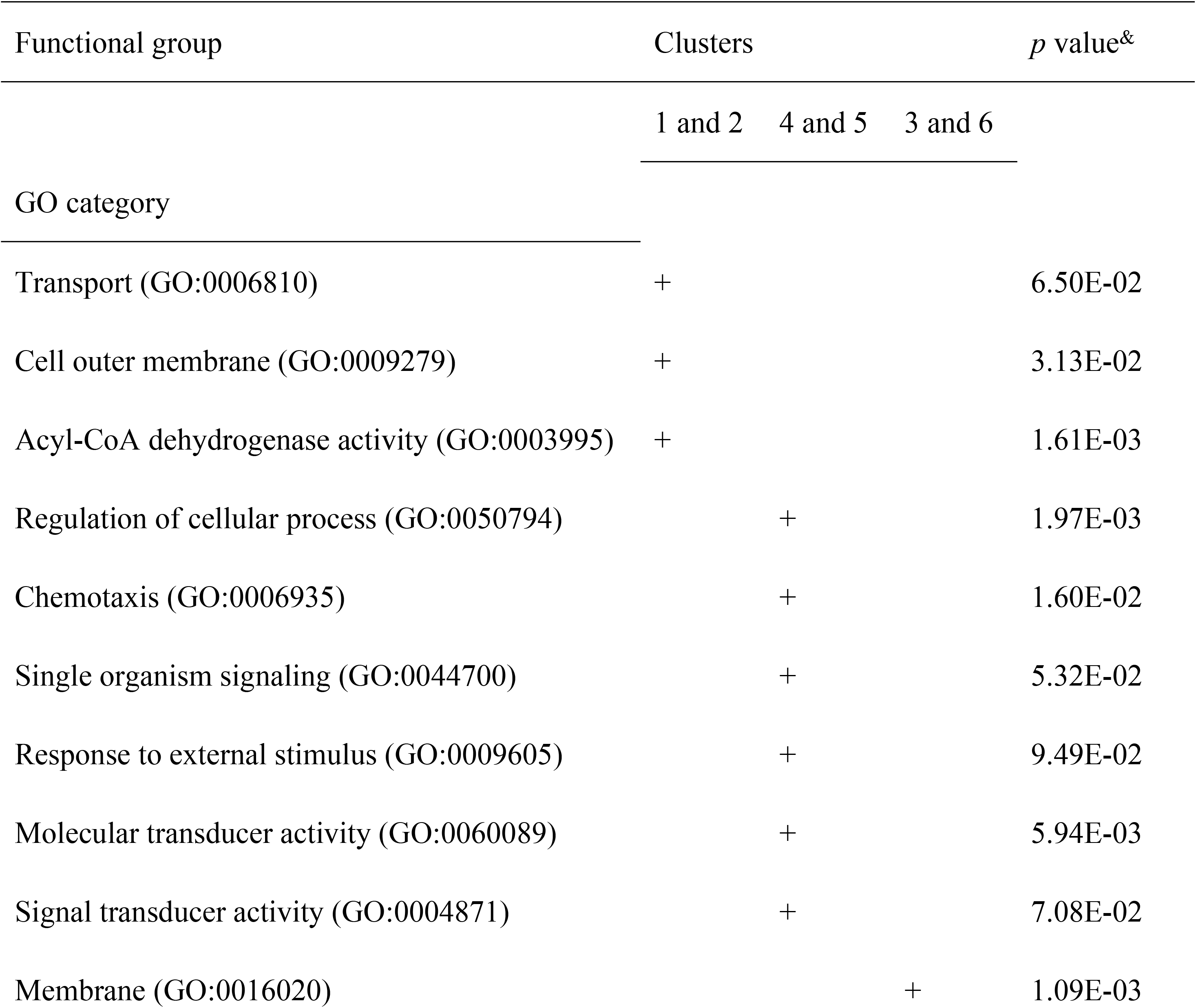
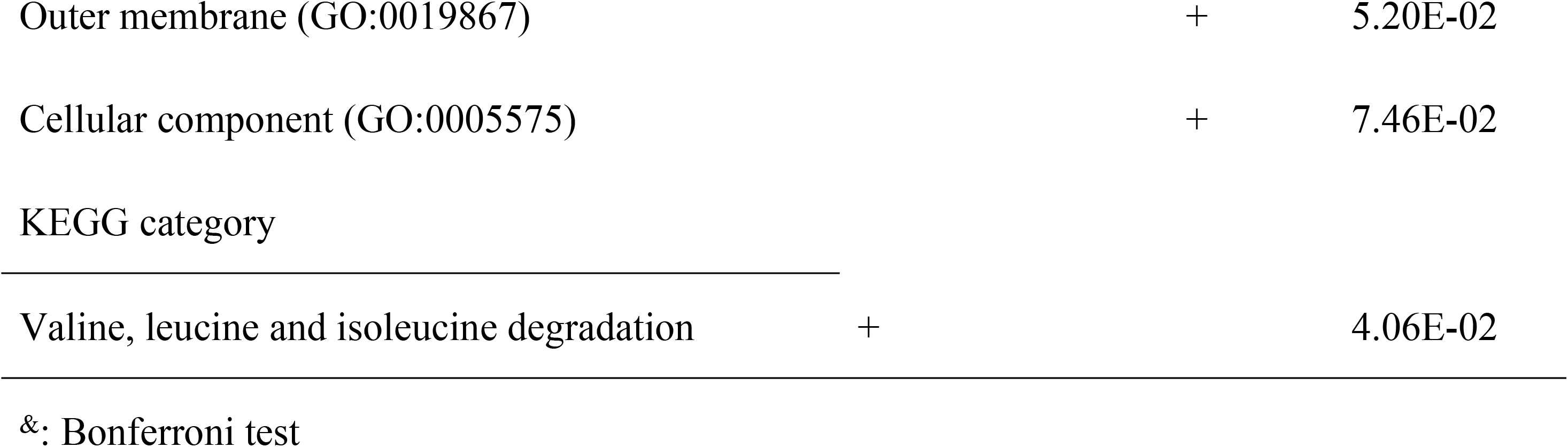
The functional groups and their significance in different gene clusters.

### Gene clusters involved in the DEGs

Since it is widely accepted that those genes forming a bacterial gene cluster are always involved in a common pathway or function [26], the DEGs located in the gene clusters we identified may play a fundamental role in oxidative stress. Based on our criterion—that at least three tandem genes must be differentially expressed to constitute a gene cluster—we identified all the possible clusters and listed them in S4 Table. Importantly, we also found a large gene cluster encoding ribosomal proteins that were up-regulated in our study (S3 Table). Apart from that result, we also found an F1F0 ATPase complex cluster that was up-regulated under oxidative stress.

### Quantitative real time PCR experiments confirms the RNA-seq dataset

The RNA-seq results were validated with a qRT-PCR analysis of four selected genes (*xoc_1643*, *xoc_1946*, *xoc_2868*, and *xoc_3249*) that encompassed a range of expression levels at 7, 15, and 45 minutes. The mRNA abundance of these transcripts at these three time-points after the H_2_O_2_ treatment followed a profile similar to that of the microarray dataset, thus validating the quality of our assay. The Pearson correlation test of the microarray against the qRT-PCR measurements yielded a correlation coefficient (R^2^=0.81, n=4), suggesting that RNA-seq dataset correlated positively and tightly with the qRT quantification (Fig 2).

**Fig 2.**
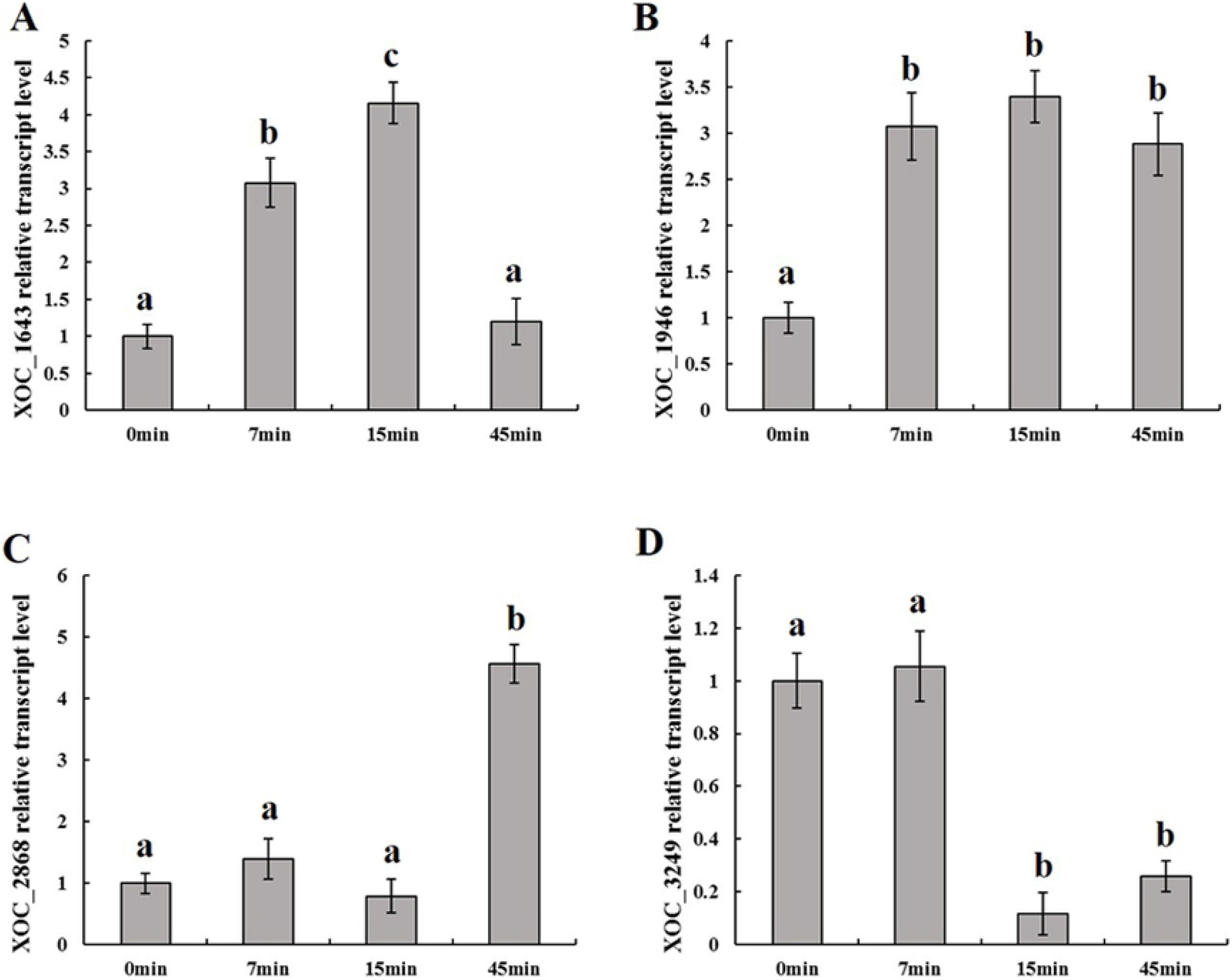
Real-time quantitative RT-PCR analysis. Transcript levels of the four candidates under oxidative stress at 7, 15, and 45 min after the H_2_O_2_ treatment. Values given are the means of five replicate measurements from a representative experiment. The experiment was repeated at least five times, and similar results were obtained. Columns with the same letters are not significantly different from each other by t-test (i.e., *P* ≥ 0.05). Error bars represent the standard deviations.

### Hypothetical gene plays some roles in the oxidative stress response in *Xoc*

Sensing, detoxification, and adaptation to oxidative stress play critical roles during successful pathogen infection and pathogenesis by *Xanthomonas* [18]. We conducted the following experiments to understand the role of these genes in *Xoc* resistance to H_2_O_2_. As Fig 3 shows, the XOC_1643 (outer membrane channel protein), XOC_3249 (membrane protein YnfA) and XOC_2868 (hypothetical protein) and mutants clearly displayed a greater sensitivity to H_2_O_2_ than did the BLS256 wild-type and complemented strains. Not surprisingly, these three mutants showed decreased pathogenicity when compared with the wild-type (Fig 4). However, the XOC_1946 (TonB-dependent receptor) mutants showed a greater resistance to H_2_O_2_ but a corresponding pathogenicity when compared with the wild-type (Fig 4). TonB-dependent receptor was proven to be involved in the transport of plant-derived molecules such as sucrose and maltodextrins in previous studies and yet delayed the disease symptom development in plants to some extent [27, 28]. Our studies show similar results with previous researches, also suggests that XOC_1946 involved in the transport of H_2_O_2_ As genes *xoc_1643*, *xoc_1946* and *xoc_3249* have proven relationships with an oxidative stress response, this result confirmed the accuracy of our RNA-seq result in this study. Interestingly, XOC_2868, a hypothetical protein, was also related with the stress response. Sequence analysis revealed that this gene has a MntR domain, thus indicating its potential role in superoxide resistance regulation [29].

**Fig 3.**
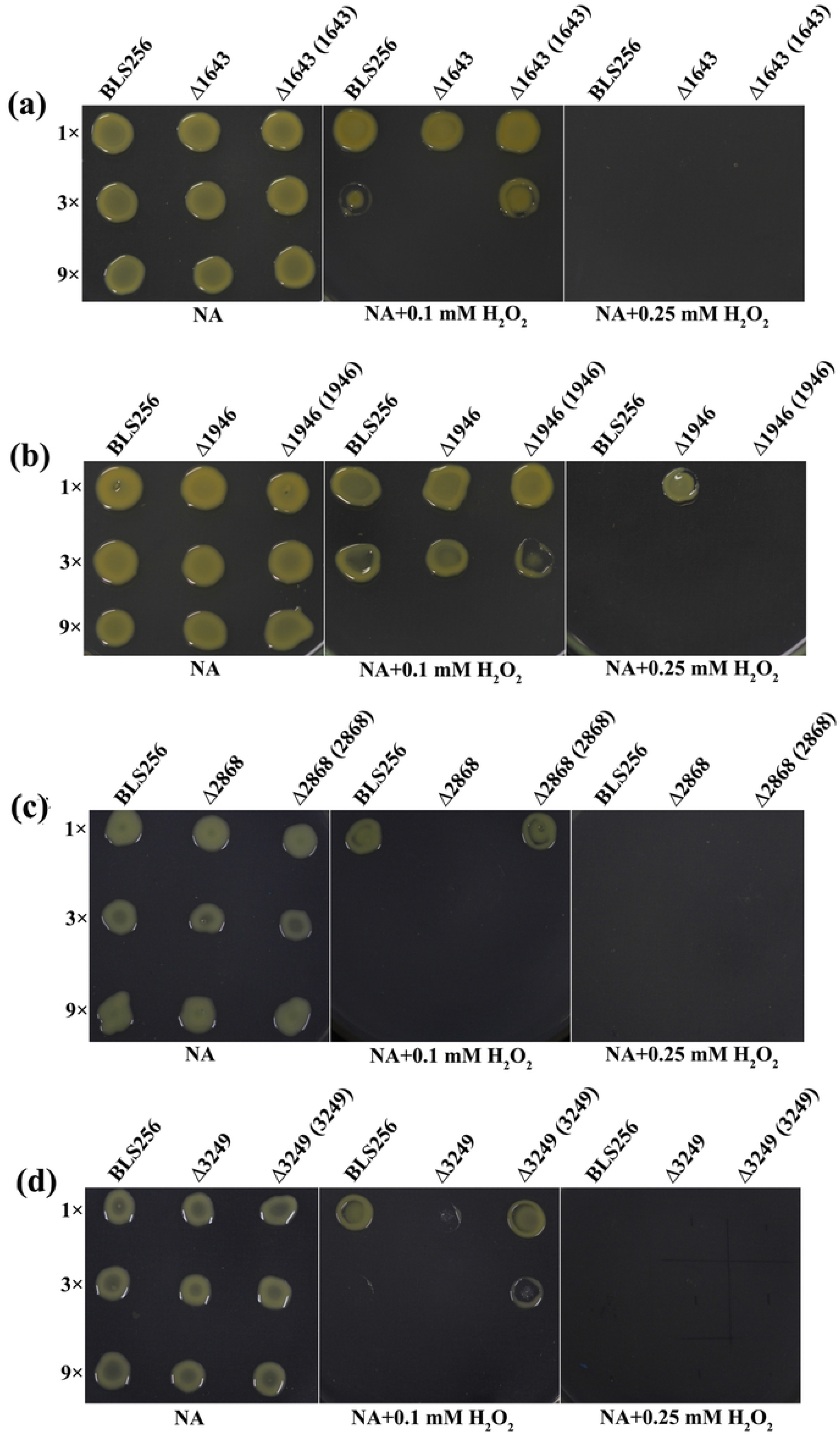
Gene mutations changed the resistance to H_2_O_2_ in *Xanthomonas oryzae* pv. *oryzicola* (*Xoc*). *Xoc* strains, including the wild-type strain BLS256, the gene deletion mutants Δ1643, Δ1946, Δ2868, and Δ3249, and their complemented strains Δ1643 (1643), Δ1946 (1946), Δ2868 (2868), and Δ3249 (3249), were grown on nutrient broth agar (NA) plates with 0 mM H_2_O_2_, 0.1 mM H_2_O_2_, or 0.25 mM H_2_O_2_. Three replicates for each treatment were used, and the experiment was repeated three times.

**Fig 4.**
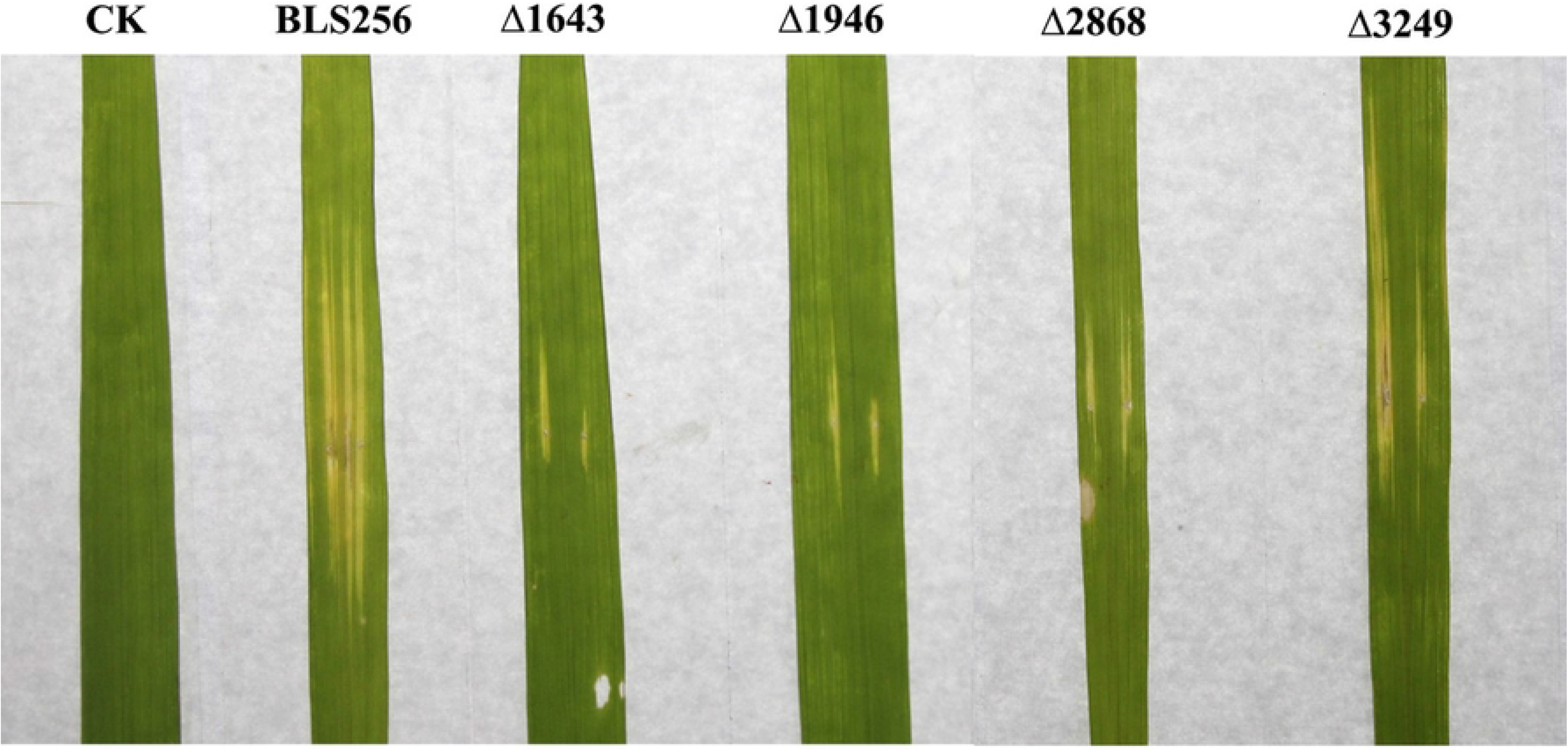
Gene mutations changed the virulence and growth of *X. oryzae* pv. *oryzae in planta*. Symptoms induced by different *X. oryzae* pv. *oryzae* strains inoculated to leaves of 2-month-old rice plants (IR24, a susceptible cultivar). Photographs were taken 14 dpi. Three replicates for each treatment were used, and the experiment was repeated three times.

### Accession numbers

All the RNA-seq data has been deposited in GEO database (https://www.ncbi.nlm.nih.gov/geo/info/) and the accession number is from SRR4533260-SRR4533267.

## Discussion

Indeed, the trigger of these proteins is reportedly involved in many different kinds of stress responses such as oxidative stress, iron stress and zinc stress [30]. Researchers also found that TBDRs are required for full virulence of *Xanthomonas campestris* pv. *campestris* to *Arabidopsis* [27]. We may infer that TBDRs may play some role in the oxidative response in *Xoc*. Not surprisingly, many reported oxidative stress-associated gene clusters were found involved, such as those for the alkyl hydroperoxide reductase, *suf* operon and *pst* operon [31, 32].

Prior research has demonstrated that H_2_O_2_ causes a slower rate of ribosomal run off, while the expression of several ribosomal proteins associated with the translation of the stress response-associated genes was increased [33]. This finding may indicate that this gene cluster contributes to the translation of oxidative stress associated genes in our *Xoc* strain.

It is well-known that the oxidative stress generated in the plant response to pathogens will decrease the intracellular pH [33]. Although direct evidence is lacking for *Xanthomonas* species, the mutants of these genes from several other bacteria showed clear growth defects under low pH, thus indicating this complex cluster is important for maintain the △pH [34].

It is now widely appreciated that a time-series transcriptome analysis can help us to better understand how organisms react to stress conditions over time [35]. Here, we set three time points corresponding to the early, middle, and late response stages. Very few genes were significantly differentially expressed in the early stage (S3 Table). Though interestingly, we did find that one gene encoding *soxR*, a redox-sensitive transcriptional activator, was the highest up-regulated gene in this time stage. In *E. coli*, *soxR* and *soxS* were shown to control the superoxide response regulon of *E. coli* [36]. Since *Xoc* lacks the homolog of *soxS*, and *soxR* is the only transcriptional regulator, we may infer that this gene triggers the *Xoc* oxidative stress response.

A comprehensive BLAST sequence analysis revealed that this MntR-like gene occurs widely in *Xoc* but not in other *Xanthomonas* species (Fig 5). Interestingly, the homologs of this gene were found to exist in many *Burkholderiaceae* family bacteria, such as *Burkholderia*, *Ralstonia*, and *Cupriavidus* (Fig 5). The phylogenetic analysis further suggested that this gene in *Xoc* might have originated from a transfer from a *Burkholderiaceae* ancestor over the course of evolutionary history, an inference with high bootstrapping support (Fig 5). Notably, many genes located adjacent to XOC_2868 are transposase (i.e., from XOC_2859 to XOC_2865). Ochman et al. suggested that HGT is mediated by bacteriophage integrases or mobile element transposases, while Keeling et al. suggested that a phylogenetic tree is the gold standard by which to identify HGT [37, 38]. The results presented here strongly suggest that *xoc_2868* is a horizontally transferred gene.

**Fig 5.**
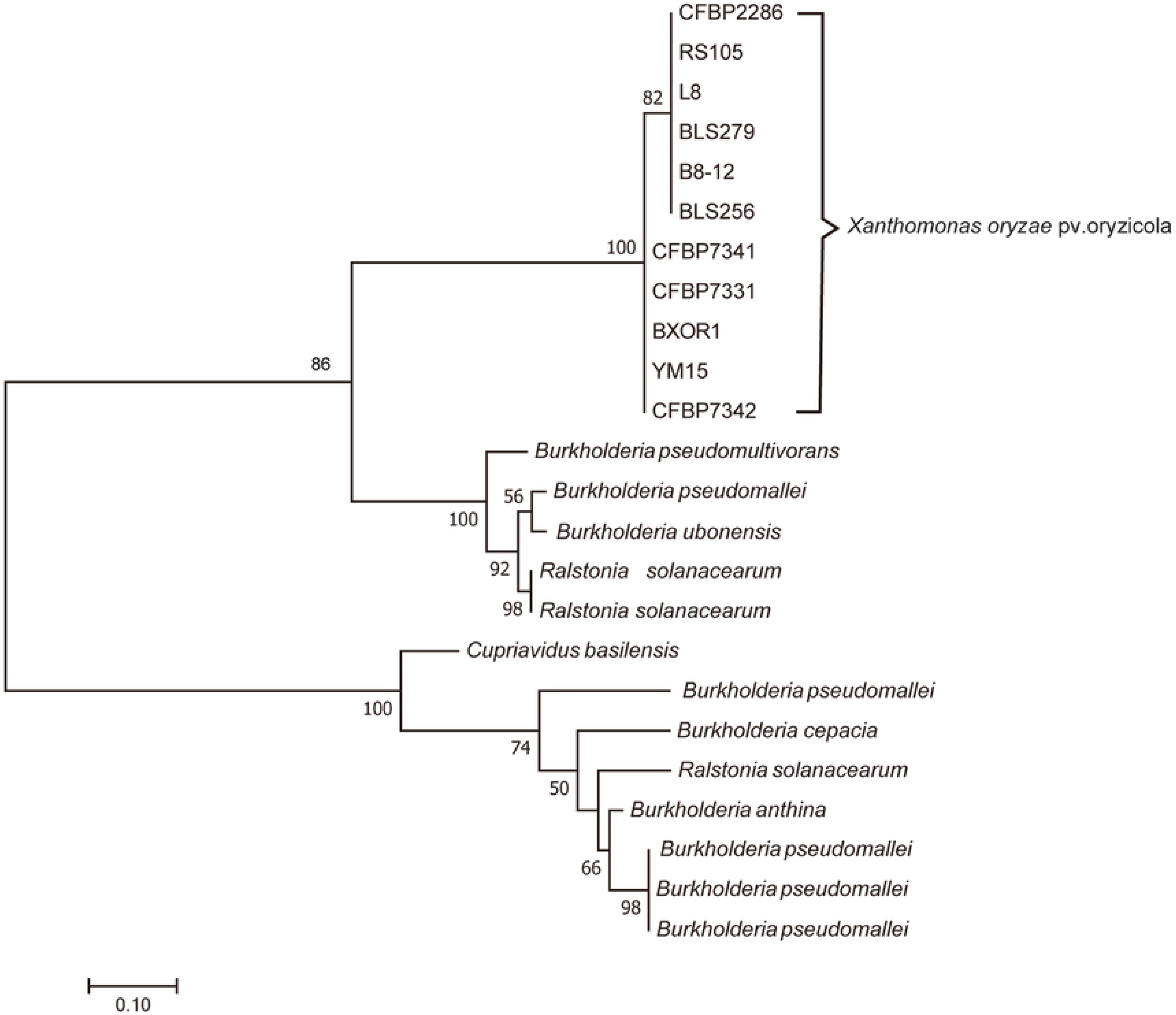
A phylogeny of the XOC_2868, a MntR-like gene. The phylogenetic tree shown was calculated using the maximum likelihood (ML) program in PhyML (Guindon *et al.*, 2010). Only the ML values ≥ 50% were shown. Bar = 0.1 substitution per site.

## Conclusions

In this research, gene expressions of *Xoc* strain BLS256 in response to a time-series H_2_O_2_ treatment have been presented by RNA-Seq analysis. In general, 7, 177, and 246 genes were differentially regulated in the early, middle, and late stages, respectively. *SoxR* gene was highly up-regulated in the early stage, indicated that this gene triggers the *Xoc* oxidative stress response. In addition, the sensitivity to H_2_O_2_ and pathogenicity of four DEGs’ mutants have been investigated, the results prove great relationships between these DEGs with an oxidative stress response. Interestingly, the results about a hypothetical protein XOC_2868 presented here strongly suggest that it is a horizontally transferred gene.

## Acknowledgements

The present study was supported by Zhejiang National Natural Science Foundation of China (LY18C140003) and National Natural Science Foundation of China (31200003).

## Supporting information

**S1 Table.** Strains and plasmids used in this study.

**S2 Table.** Primers used in this study.

**S3 Table.** Differential expressed genes in the early, middle and late stage.

**S4 Table.** Gene clusters that were differential expressed in the early (red color), middle (yellow color) and late (blue color) stages.

